# *Strongyloides* RNA-seq Browser: a web-based software platform for on-demand bioinformatics analyses of *Strongyloides* species

**DOI:** 10.1101/2021.02.18.431867

**Authors:** Astra S. Bryant, Stephanie F. DeMarco, Elissa A. Hallem

## Abstract

Soil-transmitted gastrointestinal parasitic nematodes infect approximately 1 billion people worldwide, predominantly in low-resource communities. Skin-penetrating gastrointestinal nematodes in the genus *Strongyloides* are emerging as model systems for mechanistic studies of soil-transmitted helminths due to the growing availability of functional genomics tools for these species. To facilitate future genomics studies of *Strongyloides* species, we have designed a web-based application, the *Strongyloides* RNA-seq Browser, that provides an open source, user-friendly portal for accessing and analyzing *Strongyloides* genomic expression data. Specifically, the *Strongyloides* RNA-seq Browser takes advantage of alignment-free read mapping tools and R-based transcriptomics tools to re-analyze publicly available RNA sequencing datasets from four *Strongyloides* species: *Strongyloides stercoralis, Strongyloides ratti, Strongyloides papillosus*, and *Strongyloides venezuelensis*. This application permits on-demand exploration and quantification of gene expression across life stages without requiring previous coding experience. Here, we describe this interactive application and demonstrate how it may be used by nematode researchers to conduct a standard set of bioinformatics queries.

## INTRODUCTION

Soil-transmitted gastrointestinal parasitic nematodes, including those in the genus *Strongyloides*, are a major source of disease and economic burden. *Strongyloides* species infect a range of human and animal hosts; the human parasite *Strongyloides stercoralis* is estimated to infect approximately 610 million people worldwide (Buonfrate *et al*. 2020). Similar to other intestinal parasitic nematodes, *Strongyloides* species have complex life cycles (Figure 1A) that include: parasitic females living inside a host that reproduce via asexual parthenogenesis; eggs and post-parasitic larval stages (L1-L2) that primarily exist on feces excreted from host animals; a developmentally arrested third larval stage (iL3) that actively searches for hosts to infect; and activated iL3s that have successfully entered a host animal and have resumed feeding and development (Roberts and Janovy 2005; Schafer and Skopic 2006). Unlike some intestinal parasitic nematodes such as hookworms, for *Strongyloides* species the progeny of parasitic females may develop through a limited number of free-living generations: eggs and larvae exuded from host animals develop into free-living males and females that reproduce sexually, generating post-free-living larvae that exclusively develop into iL3s (Roberts and Janovy 2005; Schafer and Skopic 2006).

**Figure 1.**
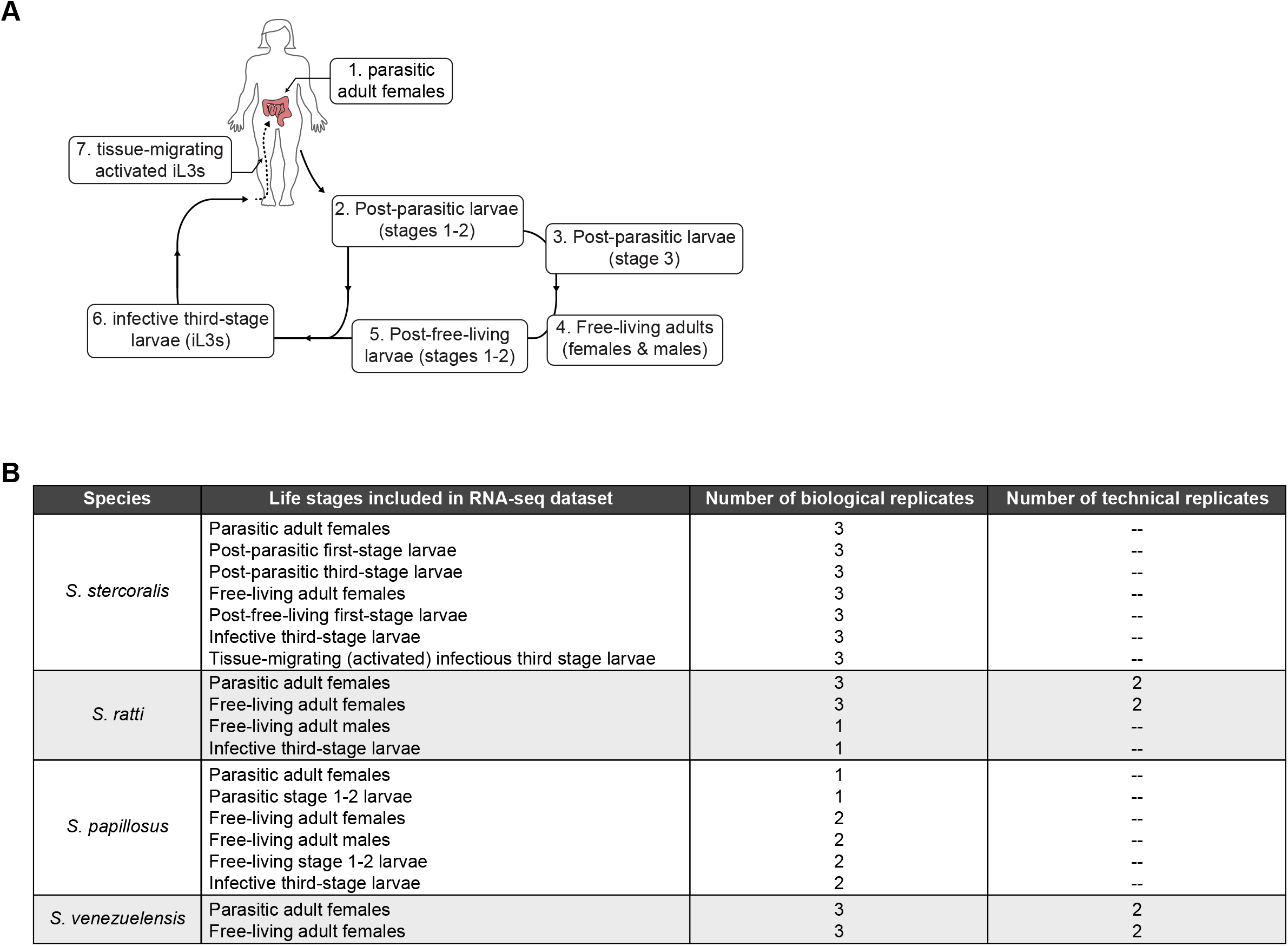
Life cycle of *Strongyloides* species. **A)** Schematic of the developmental life cycle of *Strongyloides* species, including: stages that reside within host animals (parasitic adult females and tissue-migrating, activated iL3s); free-living adults and larvae that primarily reside on feces excreted from host animals (post-parasitic larvae, free-living adult males and females, and post-free-living larvae); and developmentally arrested third-stage larvae that migrate away from host feces to actively search the environment for host animals. **B)** Life stages included in the *Strongyloides* RNA-seq Browser, by species. The number of biological and technical replicates are listed for each life stage, when available. For *S. stercoralis*, *S. ratti*, and *S. papillosus*, all publicly available samples are included in the app. For *S. venezuelensis*, data from free-living adult females and parasitic adult females are included; additional available life stages (eggs, L1, various iL3 stages) are excluded. Analyses and discussion related to this decision are available at https://github.com/HallemLab/Bryant-DeMarco-Hallem-2021.

Ongoing advances in functional genomics techniques such as transgenesis and CRISPR/Cas9-mediated mutagenesis are positioning *Strongyloides* species as genetically tractable model systems for gastrointestinal parasitic nematodes (Lok *et al*. 2017; Castelletto *et al*. 2020). The enhanced genetic tractability of *Strongyloides* species relative to other parasitic nematodes is due in large part to the availability of free-living reproductive adults, a life stage specific to the genera *Strongyloides* and *Parastrongyloides* (Castelletto *et al*. 2020). Intragonadal microinjection of exogenous DNA into *Strongyloides* free-living adult life stages can be used to generate transgenic post-free-living progeny, including infective larvae (Lok *et al*. 2017; Castelletto *et al*. 2020). Microinjection of plasmid constructs containing *Strongyloides*-specific regulatory elements permits cell-type-specific expression of a range of transgenes (Lok *et al*. 2017; Castelletto *et al*. 2020). Furthermore, the recent adaptation of CRISPR/Cas9-mediated targeted mutagenesis protocols for *Strongyloides* species allows researchers to disrupt the function of specific genes (Lok *et al*. 2017; Gang *et al*. 2017). In some *Strongyloides* species, genomic integration of transgenes has been used to establish stable transgenic lines, a necessity for mechanistic studies of free-living life stages (Lok *et al*. 2017; Castelletto *et al*. 2020). These technical advances are greatly facilitated by the fully sequenced genomes of four *Strongyloides* species: *S. stercoralis*, the rodent parasites *Strongyloides ratti* and *Strongyloides venezuelensis*, and the ruminant parasite *Strongyloides papillosus* (Hunt *et al*. 2016; Howe *et al*. 2017). High-quality reference genomes permit researchers to identify homologous genes in parasitic and free-living nematodes, while transgenesis and mutagenesis protocols allow researchers to test the functional contribution of these genes to parasitic behaviors (Gang *et al*. 2017, 2020; Bryant *et al*. 2018).

In *Strongyloides* species with fully sequenced genomes, bulk RNA sequencing datasets are also publicly available, and their analysis has provided insight into the genetic and evolutionary basis of parasitism (Stoltzfus *et al*. 2012; Hunt *et al*. 2016, 2018). However, published results generally focus on only a subset of the life stages for which RNA-seq data is available (Stoltzfus *et al*. 2012; Hunt *et al*. 2016, 2018), or are not easily adapted for quantitative comparisons (Howe *et al*. 2017). The field has lacked a user-friendly portal for accessing quantitative *Strongyloides* gene expression data and performing bioinformatics queries across life stages. Here, we present a *Strongyloides* RNA-seq Browser that permits exploration of RNA expression levels in four *Strongyloides* species and features a streamlined user interface for on-demand differential gene expression and functional enrichment analyses. We hope that this broadly accessible tool will support future studies of the genetic basis of parasitism in *Strongyloides* nematodes.

## MATERIALS AND METHODS

### Pre-processing pipeline

A kallisto::limma-voom preprocessing pipeline was implemented for each species (Figure S1).

### Data Source

Raw reads and study design files (File S1) were downloaded from the European Nucleotide Archive using the following study accession numbers: PRJEB3116 (*S. stercoralis);* PRJEB1376 and PRJEB3187 (*S. ratti*); PRJEB14543 (*S. papillosus);* and PRJDB3457 (*S. venezuelensis)*. The following reference transcriptomes were downloaded from WormBase ParaSite:

PRJEB528.WBPS14.mRNA_transcripts (*S. stercoralis)*

PRJEB125.WBPS14.mRNA_transcripts (*S. ratti)*

PRJEB525.WBPS14.mRNA_transcripts (*S. papillosus)*

PRJEB530.WBPS14.mRNA_transcripts (*S. venezuelensis)*

### Kallisto alignment and gene annotation

For each species, Kallisto was used to perform ultra-fast read mapping of raw reads to a reference transcriptome (Bray *et al*. 2016). Quality controls for raw data and Kallisto alignments were assessed and summarized using FastQC v0.11.9 and MultiQC v1.9 (Andrews 2010; Ewels *et al*. 2016). Kallisto-generated transcript abundance data was imported into R v3.6.3 using the tximport package v1.14.2; transcript read counts were generated using the lengthScaledTPM option (Soneson *et al*. 2015).

Gene annotations were imported from WormBase ParaSite via the biomaRt package v2.42.1 (Durinck *et al*. 2005, 2009). Annotation information includes: *C. elegans* homologs/percent homology, UniProKB number, InterPro terms, and Gene Ontology (GO) terms. The four *Strongyloides* species can be divided into two distinct subclades: *S. venezuelensis – S. papillosus* and *S. ratti – S. stercoralis* (Hunt *et al*. 2016, 2018). Thus, we also include gene homologs/percent homology relative to the appropriate ingroup species and a reference member of the out-group (either *S. stercoralis* or S*. papillosus*).

### Data filtering and normalization

Transcript read counts were converted to log2 counts per million (log_2_CPM) using the edgeR package v3.28.1 (Robinson *et al*. 2010), then filtered to remove transcripts with low counts (File S2). The filtering criteria varied across species as follows, depending on the number of biological replicates (Figure 1B): *S. stercoralis* and *S. venezuelensis*, 1 log_2_CPM in at least 3 samples; *S. ratti* and *S. papillosus*, 1 log2CPM in at least 1 sample.

Filtered read counts were normalized using the trimmed mean of M-values (TMM) method (Robinson and Oshlack 2010) to permit between-samples comparisons. The mean-variance relationship was modeled using a precision weights approach via the voom function in the limma package v3.42.2 (Law *et al* 2014). For *S. venezuelensis* and *S. ratti*, the RNA-seq study design included technical replicates (Figure 1B); in those cases, voom-modeled data were condensed by replacing technical replicates with their weighted average using the limma::avearrays function.

### Application overview, function, and benchmarking

Interactive app functionality was implemented using the Shiny R package v1.5.0.

### In-app gene lookup

Users may search for genes of interest using the following identifiers: gene IDs with prefixes “SSTP,” “SRAE,” “SPAL,” “SVE;” keywords matched against WormBase ParaSite gene descriptions; InterPro term; or parasite Ensembl Compara protein family names (Hunt *et al*. 2016). For *S. ratti*, users may also search for gene IDs with the prefix “WB.” Users may provide *C. elegans* gene names, which will retrieve known *Strongyloides* homologs based on WormBase ParaSite classifications.

### In-app differential gene expression

The limma package is used to conduct pairwise differential gene expression (DGE) analyses between life stages (Ritchie *et al*. 2015; Phipson *et al*. 2016). Specific contrasts are defined via user inputs. For all species, variance-stabilized, filtered, normalized log2CPM values are fit to a linear model using the limma::lmFit function. The design matrix used to fit the linear model specifies no intercept/blocking, with comparisons across life stages. To provide increased statistical power, we use empirical Bayes smoothing of gene-wise standard deviations via the limma::eBayes function (Smyth 2004). Differentially expressed genes are identified using the limma::decideTests function: *p*-values are adjusted for multiple gene-wise comparisons using the Benjamini-Hochberg false discovery rate (FDR) method. When optional correction for multiple pairwise comparisons is specified, the “global” method for multiple testing is applied; otherwise the “separate” option is used. Significantly expressed genes are classified based on an FDR of ≤0.05 and an absolute log fold change (log_2_FC) of ?1. In “Browse by Gene” mode, DGE analyses are conducted first on the entire genome; then results for user-specified genes of interest are extracted.

### Functional enrichment analysis

Gene set enrichment analysis (GSEA) is performed via the GSEA function in the clusterProfiler package v3.14.3 (Yu *et al*. 2012) using gene sets extracted from a database of parasite Ensembl Compara protein families (Hunt *et al*. 2016). On-demand GSEA is performed on gene lists rank-ordered by log_2_FC. Offline GSEA of genes contributing to PC1/PC2 identity was performed on a gene list rank-ordered by PC1 or PC2 gene scores; the two ranked lists were analyzed independently, then merged for plotting (Figure S4).

### In-app data visualization

For heatmaps of log_2_CPM gene expression, life stages are ordered using Spearman clustering and genes are ordered using Pearson clustering; clustering is performed on user-defined genes using the stats package. Heatmaps are plotted using a local copy of the ggheatmap function in the heatmaply package v1.1.1 (Galili *et al*. 2018); this copy alters the default plot margins, as well as how tick labels and plot titles are displayed. Clustering dendrograms are calculated and plotted using the dendextend package v1.4.0. Plots of individual gene expression across life stages, DGE volcano plots, and GSEA bubble plots are generated using the ggplot2 package v3.3.2 (Wickham 2009). Interactive tables are generated using the DT package version 0.14 and saved using the openxlsx package v4.2.3. Users may download any plot or table as a PDF or Excel file, respectively. When downloading DGE analysis results as an Excel file, users are offered several filtering options, including: download genes with a specific direction of differential expression *(e.g.*, only upregulated genes); download a specific percentage of differentially expressed genes (based on log_2_FC value); only download genes with selected expression types in all searched pairwise comparisons (*e.g*., only genes upregulated in iL3s relative to all other life stages).

### PCA and benchmarking analyses

For each species, principal component analysis (PCA) was run on filtered, normalized log_2_CPM data using the stats::prcomp function. For statistical comparisons of expression levels in genes exclusively found in the *Strongyloides* RNA-seq Browser relative to genes also included in benchmarking datasets, the median log2CPM expression values across biological replicates were calculated for each life stage. Next, 2-way ANOVAs (Type III) and Tukey HSD *post-hoc* tests were run using the stats package and the car package version 3.0-8.

### Data availability

The source code for species-specific preprocessing pipelines, including MultiQC reports; example analyses, including additional multivariate quantification and plots; and other supplemental materials are available at: https://github.com/HallemLab/Bryant-DeMarco-Hallem-2021

A web-hosted version of the app is available at: https://hallemlab.shinyapps.io/strongyloides_rnaseq_browser/

Due to Shinyapps.io memory constraints, computationally intensive analyses *(e.g.*, exploring gene expression for >2,000 genes) may require users to run apps locally.

App source code and deployment instructions are available at: https://github.com/HallemLab/Strongyloides_RNAseq_Browser

Supplemental figures and the following supplemental files have been uploaded to figshare. File S1 contains metadata tables for the four *Strongyloides* species included in the *Strongyloides* RNA-seq Browser. For all species, metadata information was retrieved from the European Nucleotide Archive. For *Strongyloides stercoralis*, additional protocol information was retrieved from ArrayExpress. File S2 contains low-count genes and associated log2CPM values discarded during preprocessing. File S3 contains a list of the top 10% of genes contributing to PC1 and PC2 identity for each species. Genes were extracted based on PCA variable scores using tidyverse package version 1.3.0. File S4 contains a code freeze of the *Strongyloides* RNA-seq Browser.

File S5 is an example .xlsx results file generated by the *Strongyloides* RNA-seq Browser containing the top 10% of upregulated and downregulated *S. stercoralis* genes in the iL3 versus free-living female life stages. File S6 is an output table with results of GSEA analysis of differentially expressed genes in *S. stercoralis* iL3s versus free-living females. File S7 contains a list of chemoreceptor gene IDs for all *Strongyloides* species (Wheeler *et al*. 2020; Langeland *et al*. 2020). File S8 is an example .xlsx results file generated by the *Strongyloides* RNA-seq Browser containing a differential gene expression table listing 64 *S. stercoralis* chemoreceptors that are consistently upregulated in iL3s versus all other life stages.

## RESULTS AND DISCUSSION

### Data content

The *Strongyloides* RNA-seq Browser contains bulk RNA sequencing data from four *Strongyloides* species: *S. stercoralis, S. ratti, S. papillosus*, and *S. venezuelensis*. These four species can be divided into two evolutionarily divergent subclades featuring pairs of more closely related species: *S. stercoralis – S. ratti* and *S. papillosus – S. venezuelensis* (Hunt *et al*. 2016, 2018). For each species, the number and type of developmental life stages included in the application varies based on available data (Figure 1B, File S1). We applied a preprocessing pipeline to each species in parallel (Figure S1). Key pipeline elements include: a shell script for quality control assessments and Kallisto-based alignment-free read mapping; an R Markdown (.rmd) file for conversion of abundance data to log2CPM, followed by filtering, normalization, and gene annotation; and finally, saving as static *Strongyloides* Browser input files.

For each species, we assessed the variability in expression across RNA-seq data samples by performing principal components analysis (PCA) using filtered, normalized gene expression data (Figure S2). In general, samples representing the same life stage clustered together on plots of the first two principal components for each species; biological and technical replicates of individual life stages were similar (Figure S2, Figure S3). Consistent with previous findings (Lu *et al*. 2020), we observed that for *S. stercoralis*, the first two principal components distinguish between adult and larval life stages (PC1; Figure S3) and between non-infectious and parasitic life stages (PC2; Figure S3). To test whether genes influencing PC1 and PC2 identity in *S. stercoralis* are enriched for specific parasite gene families, we performed gene set enrichment analysis on a ranked list of PC1/PC2 scores, using previously established parasite gene families (Hunt *et al*. 2016). Gene families significantly contributing to *S. stercoralis* PC1 and PC2 included multiple families encoding hypothetical proteins as well as families that were previously identified as having putative roles in parasitism: astacin-like proteins, SCP/TAP proteins, trypsin inhibitors, and acetylcholinesterases (Figure S4) (Hunt *et al*. 2016, 2018). For all species, we extracted and saved the top 10% of genes contributing to PC1 and PC2 identity (File S3).

### Software functionality

The *Strongyloides* RNA-seq Browser has two usage modes: “Browse by Life Stage” mode and “Browse by Gene” mode (Figure 2, Figure S5, File S4). In “Browse by Gene” mode, users search for gene(s) of interest by providing gene IDs or keywords that are matched against a gene annotation database. The program next offers three plotting options: a heatmap that plots the clustered expression profiles for all genes of interest across life stages, a bar plot of log2CPM expression for each gene of interest, and a table of expression values. For both modes, users next input pairwise contrasts between life stages for limma-voom-based differential gene expression (DGE) analysis. Analysis results are displayed as downloadable volcano plots and data tables (Figure S6A, File S5).

**Figure 2.**
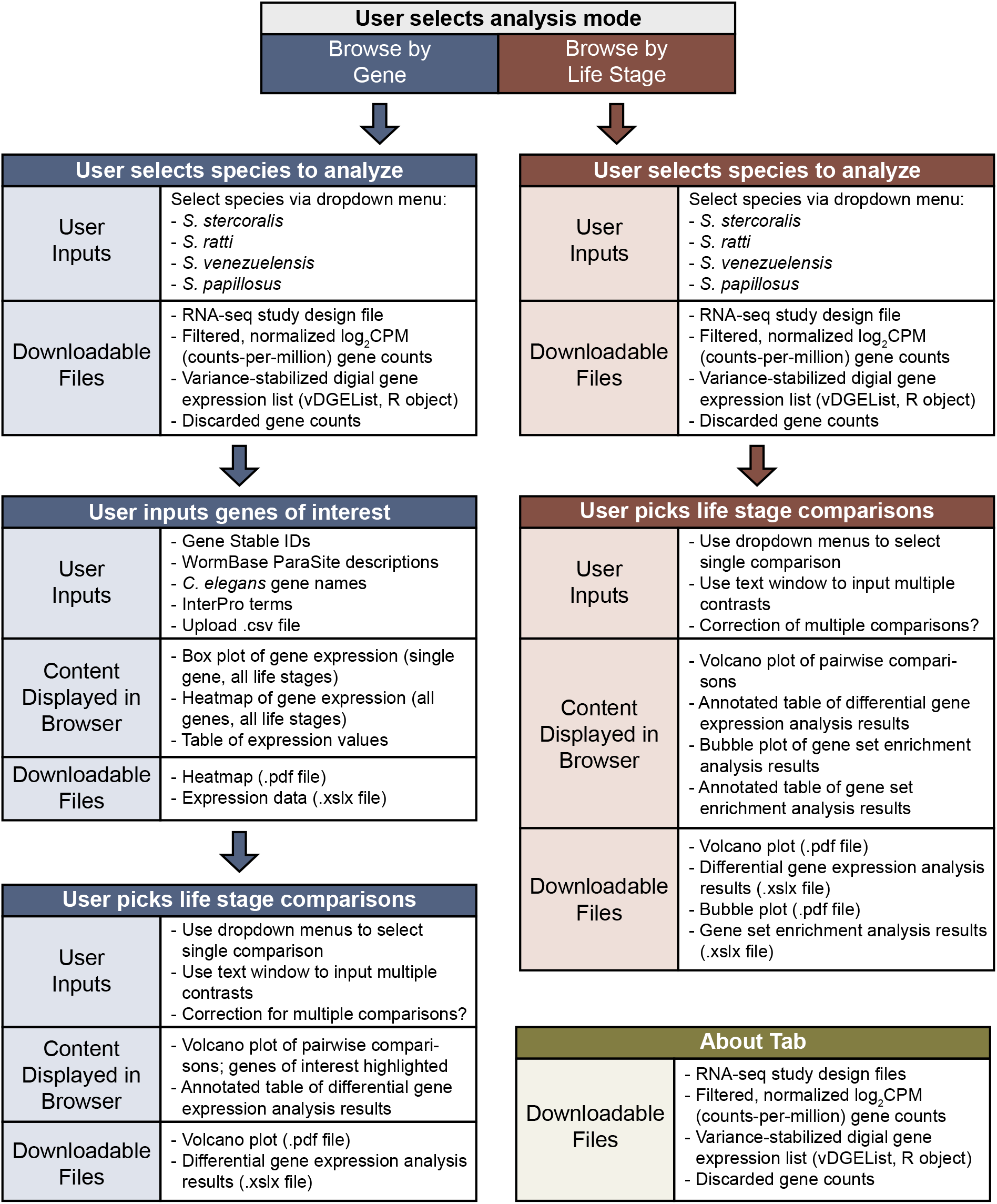
*Strongyloides* RNA-seq Browser UI overview. Overview of the *Strongyloides* RNA-seq Browser, including user inputs, content displayed in browser, and downloadable files. The application permits browsing RNA-seq data in two modes: “Browse by Life Stage” mode (red panels) and “Browse by Gene” mode (blue panels). Separate panels linked by arrows indicate the dependencies in the user experience. For example, in “Browse by Gene Mode,” users first select a species to analyze, then input genes of interest, then pick life stage comparisons. Subpanels labeled “Content Displayed in Browser” and “Downloadable Files” indicate software elements that are available after users have submitted the indicated user inputs. The user interface also includes an “About” tab that displays methods information and a menu for downloading pre-processed datafiles. For downloadable files, file formats are .csv files unless otherwise indicated.

In “Browse by Life Stage” mode, the results of DGE analyses are used to perform Gene Set Enrichment Analysis (GSEA), using an established parasite Ensembl Compara protein family database (Hunt *et al*. 2016). The GSEA analysis returns a downloadable bubble plot of enriched gene families as well as a downloadable table containing normalized gene enrichment scores that represent the degree to which the elements of the gene set are over-represented at the edges of the ranked gene list (Figure S6B, File S6).

### Benchmarking and example usage

#### Benchmarking

To assess the relative accuracy of the kallisto::limma-voom analysis pipeline, we compared *Strongyloides* RNA-seq browser data and results against previously published analyses. For *S. stercoralis* and *S. ratti*, we conducted benchmarking relative to a dataset generated via a TopHat::edgeR pipeline (Hunt *et al*. 2016). For *S. venezuelensis*, we compared results to a dataset generated via an HT-seq::cufflinks::edgeR pipeline (Hunt *et al*. 2018). In all cases, the number of genes included in the *Strongyloides* RNA-seq Browser was greater than the number of genes listed in the published datasets (Figure S7A). This discrepancy may be due to differences in the pre-processing pipelines, as well as differences in threshold cutoffs used to filter genes for inclusion in differential gene expression datasets. Indeed, genes found exclusively in the *Strongyloides* RNA-seq Browser dataset displayed lower expression levels than genes found in both datasets, for all species across life stages (Figure S7B-D). This suggests that more permissive inclusion criteria contribute to the greater size of the *Strongyloides* RNA-seq Browser dataset. To compare DGE results, we used “Browse by Life Stage” mode to perform the following pairwise comparisons: *S. stercoralis* iL3s vs. free-living females, *S. ratti* iL3s vs. free-living females, and *S. venezuelensis* parasitic females vs. free-living females. For all species, the magnitude of differential expression in individual genes was highly similar across methods (Figure 3). Thus, the *Strongyloides* RNA-seq Browser implements a differential gene expression analysis pipeline that replicates known differential expression patterns in *Strongyloides* species, while providing flexible, on-demand analysis options.

**Figure 3.**
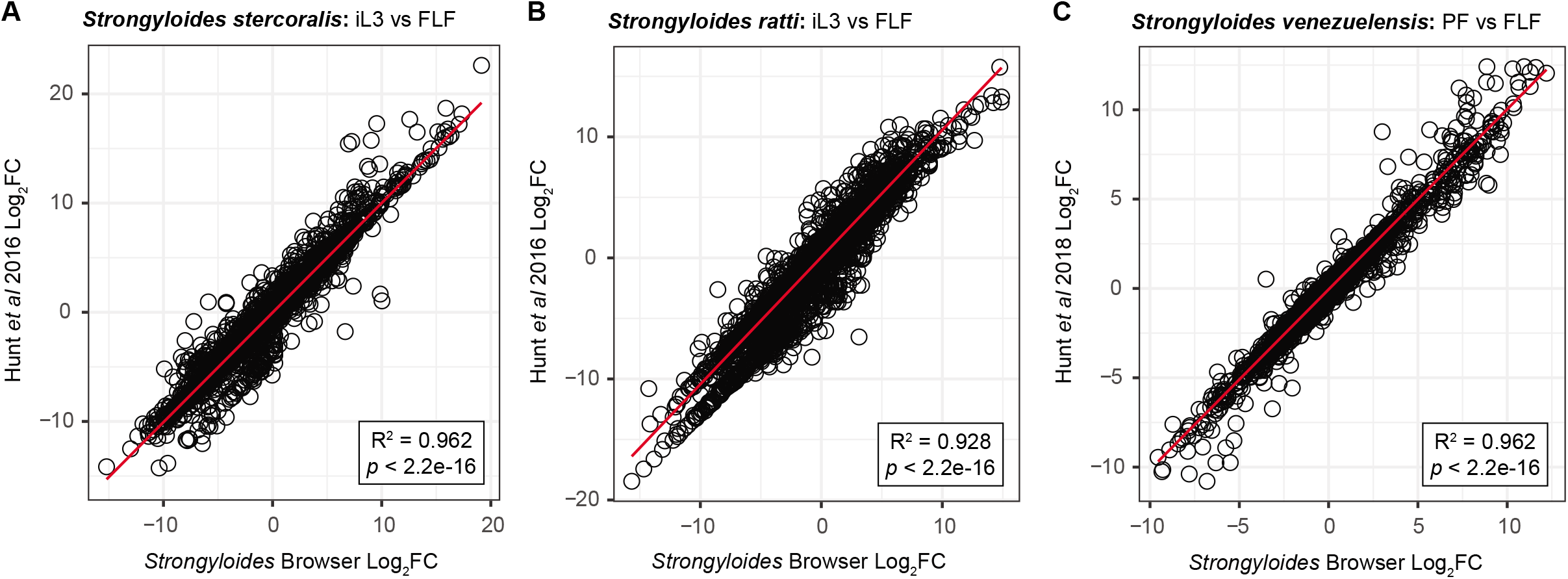
Benchmarking of selected differential expression analyses. Comparison of results generated by the *Strongyloides* RNA-seq Browser to previously published analyses in which the same datasets were analyzed using different pipelines. **A-B)** For *S. stercoralis* and *S. ratti*, iL3 versus free-living female (FLF) differential gene expression results are benchmarked against data from Hunt *et al*, 2016. **C)** For *S. venezuelensis*, parasitic female (PF) versus free-living female (FLF) differential gene expression results are benchmarked against data published in Hunt *et al*, 2018. For *S. papillosus*, log2 fold change (log2FC) values from previously published analyses are not available. For all plots, black circles represent individual genes. X-axis values are log2FC values as calculated by the *Strongyloides* RNA-seq Browser; y-axis values are previously published log2FC values. Red lines indicate a linear regression comparing previously published values to those calculated by the *Strongyloides* RNA-seq Browser. Plot inlays report R^2^ and significance values for goodness of fit. These values indicate that for individual genes, there is a high correspondence between differential gene expression as calculated by the *Strongyloides* RNA-seq Browser and previously published values. Only genes included in both the *Strongyloides* RNA-seq Browser and previously published datasets are included.

#### Analysis of chemoreceptor genes

Next, we applied the *Strongyloides* RNA-seq Browser to a published dataset of *S. stercoralis* chemoreceptor genes (Wheeler *et al*. 2020). In parasitic nematodes, chemosensation plays an important role in driving life-stage-specific, ethologically relevant behaviors such as host seeking and host invasion (Bryant and Hallem 2018; Banerjee and Hallem 2020; Wheeler *et al*. 2020). For example, *S. stercoralis* iL3s respond robustly to a number of human-associated odorants, and have distinct olfactory preferences from those of other life stages (Castelletto *et al*. 2014; Gang *et al*. 2020). Many nematode chemoreceptors are G protein-coupled receptors (GPCRs), and nematode genomes encode large numbers of predicted chemoreceptor GPCRs (Bargmann 2006; Wheeler *et al*. 2020; Langeland *et al*. 2020). Putative chemosensory GPCR genes are highly divergent across nematode species (Wheeler *et al*. 2020), leading to the functional diversity that likely underlies species-specific differences in chemosensory behaviors (Hallem *et al*. 2011; Dillman *et al*. 2012; Castelletto *et al*. 2014; Lee *et al*. 2016; Gang *et al*. 2020). We asked whether *Strongyloides* putative chemoreceptor genes displayed life-stage-specific expression patterns that may underlie life-stage-specific behaviors.

We analyzed 167 *S. stercoralis* chemoreceptor genes using “Browse by Gene” mode (File S7). Clustering analysis of log2CPM expression of *S. stercoralis* chemoreceptors across life stages revealed that expression patterns in iL3s and activated iL3s are highly similar, compared to all other life stages (Figure 4A). We next used the browser to perform pairwise comparisons between *S. stercoralis* chemoreceptor expression in iL3s versus every other life stage; these comparisons confirmed that the expression of most chemoreceptor genes is significantly higher in iL3s (log2FC ≥ 1, FDR ≤ 0.05) (Figure 4B-C). Furthermore, we identified 64 chemoreceptor genes that are selectively upregulated in iL3s relative to all other life stages (File S8). These results agree with previous observations that key elements of the chemosensory signal transduction pathway that are located downstream of GPCRs are upregulated in iL3s and that generally, genes upregulated in iL3s are associated with sensing the environment (Stoltzfus *et al*. 2012; Hunt *et al*. 2016).

**Figure 4.**
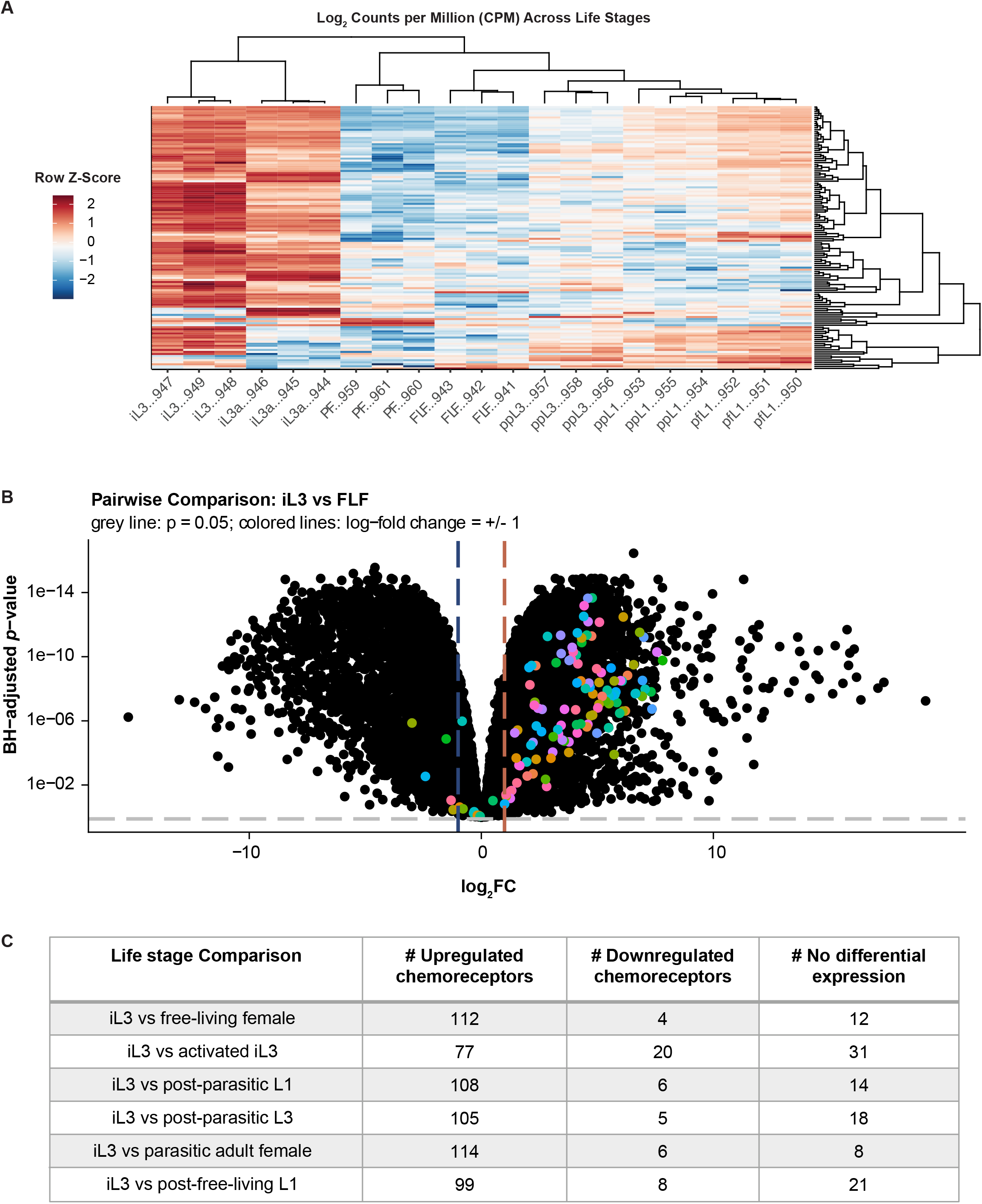
Chemoreceptor gene expression is upregulated in *S. stercoralis* infective larvae. **A)** Heatmap of log2 counts per million (CPM) expression of 128 *S. stercoralis* chemoreceptor genes across developmental life stages. Columns represent RNA-seq samples and are labeled with the appropriate life stage abbreviation and the last 3 digits of the sample ID number. Column ordering and dendrogram reflects results of Spearman clustering; row order and dendrogram reflects results of Pearson clustering. Heatmap color scale is based on a row Z-score calculated by centering and row-scaling the log2CPM values for each gene. Life stage abbreviations are as follows: infective third-stage larvae (iL3), activated iL3s (iL3a), parasitic adult females (PF), free-living females (FLF), post-parasitic first-stage larvae (ppL1), post-parasitic third-stage larvae (ppL3), post-free-living first-stage larvae (pfL1). **B)** Volcano plot of differential gene expression between iL3s and free-living females (FLF). Positive log2FC values indicate enrichment in iL3s relative to free-living females; negative log2FC values indicated enrichment in free-living females relative to iL3s. Black dots are values for all *S. stercoralis* genes; colored dots indicate chemoreceptor genes. Grey line indicates Benjamini-Hochberg-adjusted *p-*value of 0.05, colored lines indicate log_2_FC = 1. **C)** Summary of differentially expressed chemoreceptor counts in iL3s versus other life stages. Genes were categorized as significantly upregulated or downregulated if they displayed an absolute log_2_FC of ≥1 and a false-discovery-rate-corrected *p*-value of ≤0.05.

Interestingly, although most GPCRs are downregulated in non-iL3 life stages, *S. stercoralis* free-living adults display robust chemosensory behaviors, including broad attraction to host-associated odorants (Castelletto *et al*. 2014; Gang *et al*. 2020). Furthermore, despite profound increases in GPCR expression levels in *S. stercoralis* iL3s, the iL3s appear to be attracted to fewer host-associated odorants than free-living adults, a change that appears to be dominated by a lack of attraction to fecal odorants in iL3s (Gang *et al*. 2020). How the widespread upregulation of GPCRs in iL3s relates to the more selective olfactory preferences of iL3s for host odorants is not yet clear. One intriguing possibility is that the GPCR genes upregulated in iL3s may act redundantly, reinforcing iL3 attraction to a limited number of host odorants in order to ensure successful detection of host animals. Alternatively, iL3-upregulated GPCRs may confer responses to host-specific odorants that have not yet been tested but that selectively contribute to iL3-specific behaviors such as skin penetration and host invasion. In the future, characterizing the function of these GPCRs may provide critical mechanistic insight into the parasitic behaviors of *Strongyloides* nematodes.

In summary, the *Strongyloides* RNA-seq Browser prioritizes a user experience that provides access to *Strongyloides* genomics data and bioinformatics analyses without requiring extensive previous coding experience. The open-source code supports future expansion: additional bioinformatics analyses can be included as requested, and the bulk RNA-seq processing and analysis pipelines may be easily reused as additional data becomes publicly available. Ultimately, we hope that the *Strongyloides* RNA-seq Browser will facilitate future genetic studies of *Strongyloides* species by acting as a user-friendly resource for researchers seeking to understand the functional roles of specific genes and gene families in parasitic nematodes.

## Acknowledgements

The authors would like to gratefully acknowledge Dr. Dan Beiting and all others involved in the DIYTranscriptomics online course (DIYTranscriptomics.com). The authors would also like to thank Dr. Vicky Hunt for providing assistance with the published RNA-seq datasets, and Dr. Michelle Castelletto for useful discussion.

